# Statistical Correction of the Winner’s Curse Explains Replication Variability in Quantitative Trait Genome-Wide Association Studies

**DOI:** 10.1101/104786

**Authors:** Cameron Palmer, Itsik Pe’er

## Abstract

Genome-wide association studies (GWAS) have identified hundreds of SNPs responsible for variation in human quantitative traits. However, genome-wide-significant associations often fail to replicate across independent cohorts, in apparent inconsistency with their apparent strong effects in discovery cohorts. This limited success of replication raises pervasive questions about the utility of the GWAS field. We identify all 332 studies of quantitative traits from the NHGRI-EBI GWAS Database with attempted replication. We find that the majority of studies provide insufficient data to evaluate replication rates. The remaining papers replicate significantly worse than expected (*p* < 10^−14^), even when adjusting for regression-to-the-mean of effect size between discovery- and replication-cohorts termed the Winner’s Curse (*p* < 10^−16^). We show this is due in part to misreporting replication cohort-size as a maximum number, rather than per-locus one. In 39 studies accurately reporting per-locus cohort-size for attempted replication of 707 loci in samples with similar ancestry, replication rate matched expectation (predicted 458, observed 457, *p* = 0.94). In contrast, ancestry differences between replication and discovery (13 studies, 385 loci) cause the most highly-powered decile of loci to replicate worse than expected, due to difference in linkage disequilibrium.

**Author Summary:** The majority of associations between common genetic variation and human traits come from genome-wide association studies, which have analyzed millions of single-nucleotide polymorphisms in millions of samples. These kinds of studies pose serious statistical challenges to discovering new associations. Finite resources restrict the number of candidate associations that can brought forward into validation samples, introducing the need for a significance threshold. This threshold creates a phenomenon called the Winner’s Curse, in which candidate associations close to the discovery threshold are more likely to have biased overestimates of the variant’s true association in the sampled population. We survey all human quantitative trait association studies that validated at least one signal. We find the majority of these studies do not publish sufficient information to actually support their claims of replication. For studies that did, we computationally correct the Winner’s Curse and evaluate replication performance. While all variants combined replicate significantly less than expected, we find that the subset of studies that (1) perform both discovery and replication in samples of the same ancestry; and (2) report accurate per-variant sample sizes, replicate as expected. This study provides strong, rigorous evidence for the broad reliability of genome-wide association studies. We furthermore provide a model for more efficient selection of variants as candidates for replication, as selecting variants using cursed discovery data enriches for variants with little real evidence for trait association.

## Introduction

Genome-wide association studies (GWAS) have identified thousands of genetic variants associated with complex human traits [1]. GWAS are most commonly two-stage designs, with a discovery study followed up by (possibly several) internal replication studies on independent samples. Due to the number of variants tested in the typical association study, replication is only attempted for a small fraction of the discovered variants exceeding a p-value threshold adjusted for 10^6^ independent tests. The tradeoff between study power per-variant and resources, along with the strategy of testing millions of variants for association, leads to study designs where many associated variants of low effect size [2] are underpowered to be detected.

The Winner’s Curse (WC) is the systematic overestimation of effects ascertained by thresholding. This phenomenon is induced by ascertainment of the most significant GWAS signals for reporting: introducing a threshold on statistical significance means that the selected set of signals will preferentially contain variants whose effects are overestimated in a particular study sample due to chance noise (S1 Fig). This tendency of studies to overestimate their association with a phenotype in the discovery cohort might cause them to replicate at an unexpectedly low rate, increasing the apparent unreliability of results from the field. This paper relies on computationally correcting this biased overestimate of effect size, in order to produce accurate estimates of the chances for replication.

Several models for directly estimating bias in effect estimates have been developed. Parametric models, based predominantly on the theory established in [3], generate a maximum likelihood estimation of the effect estimate based on the impact of introducing a p-value threshold into the reported list of variants; thus, test statistics close to the threshold tend to be biased more severely than those more substantially exceeding the threshold. Alternatively, nonparametric bootstrap correction of the Winner’s Curse using individual-level genetic data [4] has been implemented. Evaluation of these models for binary [5,6] and quantitative [7] traits has been limited to simulations and a small number of studies, without establishing the importance of WC-correction to GWAS study design.

Further complicating matters, there is no single accepted standard for successful internal replication of a variant in a GWAS. Across the GWAS considered in this study we have observed several definitions of replication. The variability of these definitions leads to differing standards of “replicating signal” in the literature, and complicates an evaluation of replicability across the field.

Variants found to be trait-associated in GWAS are not necessarily causal [8], due to linkage disequilibrium (LD) between common variants. Causal variants are expected to replicate, whereas significantly-associated noncausal variants will only replicate if they remain linked to a causal variant in a replication study. The predicted rate of replication for noncausal variants is not trivial, as in general the causal variant in a locus is unknown and may not be assayed in the study. In particular, more GWAS now attempt discovery and replication in samples of distinct ancestries, which are expected to have substantially different LD patterns across much of the genome. Moreover, even when LD between a hidden causal variant and its observed proxy are comparable across replication and discovery, there remains an open question as to whether, and in what contexts, SNPs are expected to have comparable effect in different ancestral backgrounds; existing work, in particular using the same database from this study [9], has provided inconclusive results that may be confounded by both the Winner’s Curse and a preponderance of false positive variants.

In this paper we seek to evaluate the replicability of SNPs in genome-wide association studies across the field of human quantitative trait genetics. We specifically consider quantitative trait studies as they are underrepresented amongst theoretical work for correcting the Winner’s Curse, and represent a meaningful subset of the field (33% of papers considered) that is still sufficiently small that we may feasibly evaluate all existing studies. The NHGRI-EBI GWAS Catalog [1, 10] provides a reasonably complete database of publications claiming to report genome-wide significant associations between variants and human traits. We use this catalog as a tool to identify the vast majority of papers in the field. Using only summary data reported in these papers, we modeled the Winner’s Curse in all papers providing enough information to actually support their claims of replication. We recomputed their replication rates according to the nominal and Bonferroni standards of replication, thus introducing a standardized regime to make generalizations about replication efficiency across all studies. Together, we obtain reliable metrics to evaluate the state of human quantitative trait genetics as a reproducible scientific domain.

## Results

### Paper Quality Control

We considered all 332 GWAS papers for quantitative-traits in the database [1, 10] from journals we deemed pertinent to human genetics (see Table 1, S1 Table) that attempted replication of discovered variants. We filtered this pool, requiring study design of strict thresholding, reports of data needed to calculate bias in effect sizes [3], and related consistency criteria (see Methods, S2 Table). This reduced the pool to *k* = 100 post-QC papers (30%) for analysis.

The above counts consider each paper as a functional unit. In some cases, a single paper will publish multiple GWAS: that is, multiple phenotypes will be analyzed in the same paper. The 100 papers passing QC correspond to 134 “studies,” with 79 papers containing only a single study, and the remainder having fewer than 6 studies each. As these additional studies typically contribute a very small number of variants to our analysis, we proceed with the paper count as a more honest reflection of the scope of our analysis.

**Table 1.** Distribution of papers across journals, for journals that had *at least one* article with sufficient information for analysis. The full distribution of all journals analyzed in the study, including those with all papers excluded, is in S1 Table.

|  | Analyzed | Excluded |
| --- | --- | --- |
| Am J Hum Genet | 8 | 23 |
| Am J Med Genet B Neuropsychiatr Genet | 2 | 2 |
| BMC Med Genet | 3 | 3 |
| Circ Cardiovasc Genet | 4 | 12 |
| Front Genet | 1 | 0 |
| Gene | 1 | 1 |
| Genet Epidemiol | 1 | 1 |
| Hum Genet | 3 | 7 |
| Hum Mol Genet | 24 | 48 |
| J Med Genet | 1 | 5 |
| Nat Genet | 26 | 48 |
| PLoS Genet | 18 | 31 |
| PLoS One | 6 | 21 |
| Science | 2 | 2 |
| Total: | 100 | 204 |

### Paper Characteristics

The sum of discovery sample sizes across all analyzed papers reaches approximately 1.8 million non-unique individuals. The majority (88%) of this cumulative count have European ancestry, framing the analysis in the context of this group. This 6.7-fold over representation of European ancestry is part of uneven sampling of world populations in GWAS (Table 2).

**Table 2.** Ancestry distribution of samples included in GWAS. Rows are as follows: (1) “Totals”: number of samples of a given ancestry in analyzed papers, with redundancy between studies published multiple times; (2) “Rate in GWAS”: percentage of total samples considered that were of this ancestry; (3) “Rate in Population”: percentage of world’s population that is of this ancestry; (4) “Enrichment in GWAS”: relative over (or under) representation of ancestry in GWAS relative to its rate in the world. Ancestry labels are approximations with the standard correspondences to HapMap2 reference samples (European = CEU, East Asian = JPT+CHB, African = YRI); here, “African American” denotes samples reported with that nomenclature, which typically corresponds to 80:20 admixture between ancestral sub-Saharan African and Western European genetics [11]. All of these equivalences are oversimplifications but correspond to assumptions widely used in the field. Counts are computed from totals across all papers analyzed in this study, not adjusting for duplicate uses of the same datasets across multiple studies. Total sample sizes are maximum counts of samples assuming no per-genotype missingness is present. The totals are rounded to the nearest integer as several imputed studies reported nonintegral sample sizes. Row 3 percentages in world population are approximations based on demographic data from 2014-2015 [12, 13].

|  | European | East Asian | African | African American |
| --- | --- | --- | --- | --- |
| Totals | 1601628 | 135472 | 1226 | 80006 |
| Rate in GWAS (percent) | 88.08 | 7.45 | 0.07 | 4.4 |
| Rate in Population (percent) | 13.3 | 59.8 | 13.1 | 0.5 |
| Enrichment in GWAS (percent) | 670.8 | 12.5 | 0.51 | 821.6 |

The tally of variants these papers attempted to replicate lists 2691 non-unique variants, each passing the corresponding paper-specific p-value threshold in its discovery cohort. Many of these papers include linked variants on this list, introducing partial redundancies. We filtered dependent variants (Online Methods) to obtain 1652 loci for analysis, independent within each paper.

### Replication Rates, by Paper

At a nominal threshold *α* = 0.05, we observe 793/1652 independent loci to replicate (48%) across 100 papers. Based on the raw effect sizes reported in the discovery cohort, we would have expected 1498 loci to replicate (90.7%), significantly more than observed (two-tailed Poisson binomial *p* = 4.2 · 10^−15^). Statistical correction of WC leads to a prediction of 888 replicated loci (53.8%), 7-fold closer but still significantly more than observed (*p* < 3 · 10^−16^). Replacing the nominal threshold by Bonferroni-adjusted thresholds 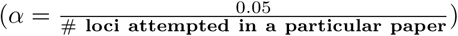, we observe 519 replicated loci (31.4%), significantly different than both raw (*p* = 3.3 · 10^−14^) and WC-corrected (*p* = 9.0 · 10^−15^) replication predictions of 1235 (74.8%) and 610 (36.9%) loci, respectively.

Predicting WC-corrected replication rates per paper (Poisson binomial distribution), we observe excess of papers both over- and under-performing their respective expectations (Fig 1A). This excess significantly correlates with publication venue (Fig 1B). Specifically, papers in higher impact journals tended to over-replicate, consistent with publication bias [14–16] (Discussion).

**Fig 1.**
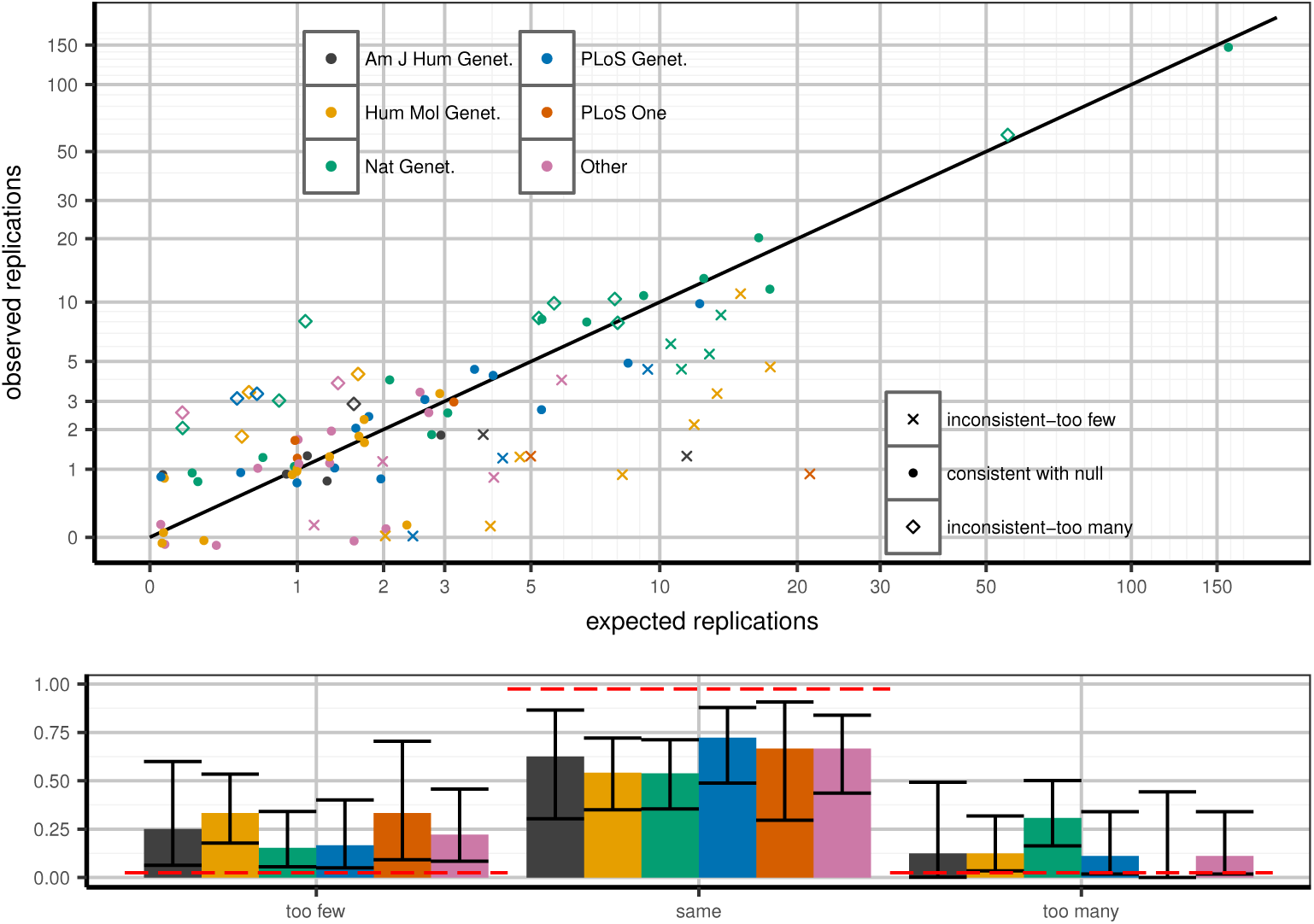
Expected and observed replication rate per publication, stratified by journal. Top panel (A): predicted versus expected replication for each paper. Each paper is flagged as being within 95% confidence of predicted replication rate under WC-corrected model (dots), greater (diamonds) or lower (Xs) than expectation. X-axis: predicted number of replications in a given paper, calculated as the sum across all loci of power to replicate based on WC-corrected discovery effect estimates. Y-axis: observed (jittered integer) number of replications in the paper. Colors correspond to journals. Replication is defined as a one-tailed replication p-value surpassing a per-paper Bonferroni threshold: 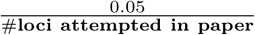. Confidence intervals defined as 95% confidence according to Poisson binomial draws from the WC-corrected power distribution. Bottom panel (B): distinct behaviors in journals depending on which set of papers is considered. Clusters correspond to paper quality (point shapes) from top panel; confidence intervals are 95% confidence intervals from the binomial distribution. Red lines are expected bar heights assuming that the observed paper data correspond to the WC-corrected model.

### Sample Size and Ancestry Explain Replication Inconsistency

Few papers (*k* = 13) discovered variants in one continental ancestry and attempted replication in another. This study design may hurt replication beyond WC due to population-specific effects, including linkage disequilibrium. Most (48/87) remaining papers reported single sample size *N* for replication across all attempted variants, not reflecting different fractions of missing data for each variant. Note that this includes genotypes missing from association analysis, rather than unmeasured genotypes whose analysis was conducted within the study, even if through imputation. In particular, studies conducting meta-analysis may only obtain variant data from a subset of their contributing cohorts, leading to large discrepancies in effective sample size per locus. This exaggerated replication sample size overestimates power to replicate and thus inflates predicted replication rate.

The remaining 39 papers with 707 discovered loci both maintained continental ancestry across discovery and replication while also correctly reporting per-locus *N.* At nominal threshold, 457 loci (64.6%) replicate, consistent (Poisson binomial *p* = 0.94) with the WC-corrected prediction of 458 loci (64.8%). Considering instead the more stringent Bonferroni correction, observed replication of 304 loci (43%) was also consistent (*p* = 0.14) with the 316 expected (44.7%). In both cases, predicting replication without WC-correction fails (all *p* < 10^−14^). Considering all thresholds across these papers, WC-correction significantly improved sensitivity over raw discovery estimates (ROC AUC 0.785 vs. 0.582, DeLong two-tailed *p* < 2 · 10^−16^; see S4 Fig). We thus hereafter consider only WC-corrected estimates.

The improved fit amongst these 39 remaining papers is not explained by reduction in power to reject fit: fit is more improved than chance expectation (based on simulations on subsets of variants with matched power to observed; nominal replication, *p* < 0.001; Bonferroni replication, *p* < 0.001). Furthermore, both *N* and ancestry filters are required for good model fit (see S6 Fig and S7 Fig).

We further tested the importance of per-locus sample size reporting by repeating the replication rate analysis on 39 papers/707 loci with correct ancestry and sample size, but instead using the maximum available sample size for each study. Correcting the Winner’s Curse using these aberrant sample sizes, the predicted rates of replication are no longer consistent with observed data (nominal replication: *p* = 0.0495; Bonferroni replication: *p* = 0.000202). These results further support the conclusion that correct per-locus sample size reporting is essential for both accurate Winner’s Curse correction and verifiable replication reporting.

### Replication Efficiency by Strength of Association

We next investigated the relationship between the strength of a signal and its replication rate. We partitioned all loci across all 100 papers into deciles according to their observed replication p-value. We then used each variant’s power to replicate within its study to predict the replication rate within each decile. Note that, as we used deciles here, the observed and expected values should both be 10%, within confidence bounds and rounding error.

Across all variants, the predicted replication rate per bin was not significantly different from 10%, as expected, with the notable exception of the highest decile: the strongest signals tended to replicate significantly less than predicted (see Fig 2). This deviation primarily explains why the entire partition into deciles was significantly different than expected (*χ*^2^ goodness of fit *p* < 10^−4^). As before, when restricting analysis to same-ancestry replication and reporting per-locus *N* (see S8-S11 Fig for other subsets), replication rates became consistent with prediction, both jointly across all decile bins (*p* = 0.67) as well as within each (S3 Table). Again, this is not simply lack of power to reject fit: the reduction in significance is beyond random expectation (*p* < 0.01). Several other partitions of the data approached good fit (S3 Table), but no more than was expected due to reduction in power (all *p* > 0.05).

**Fig 2.**
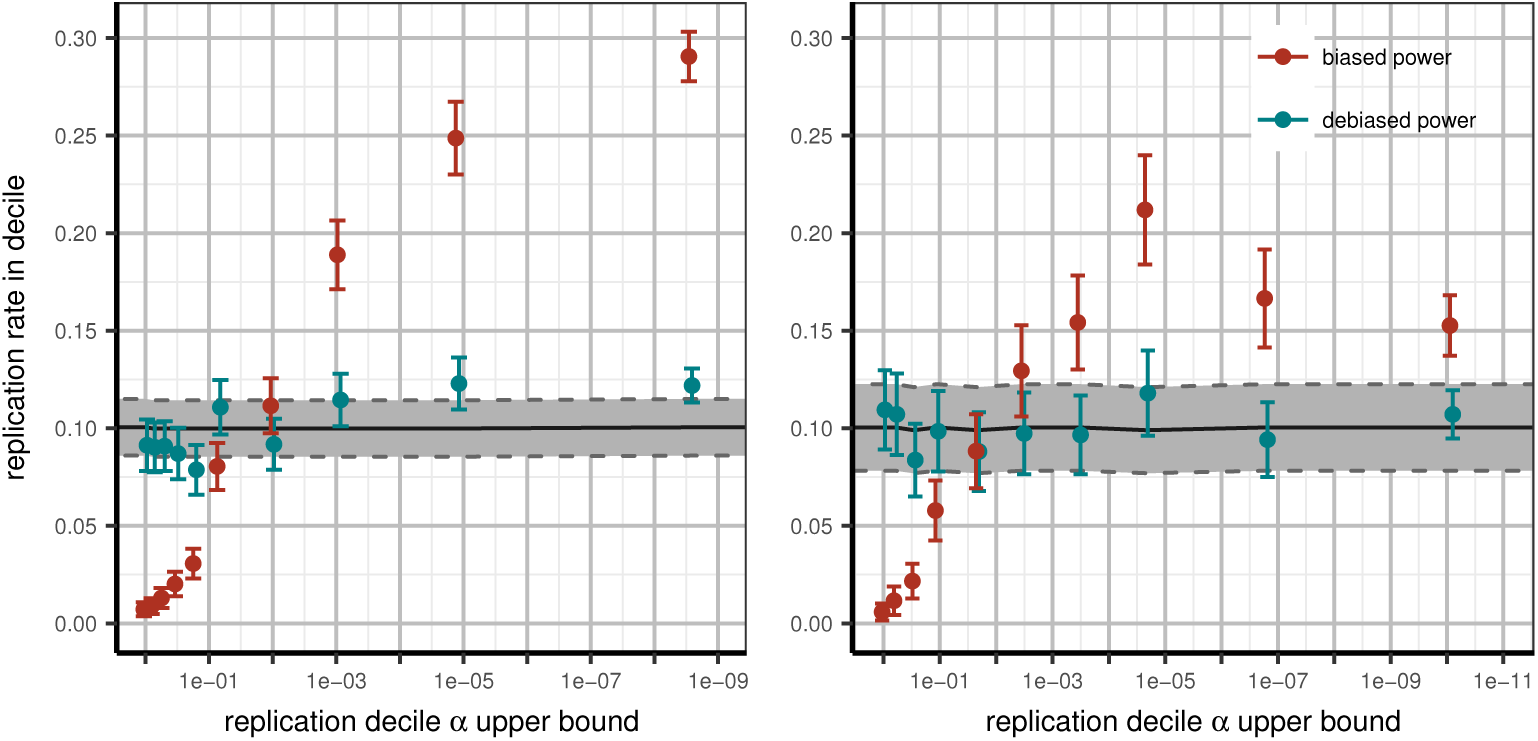
Expected and observed rates of replication in replication deciles. All variants are sorted by replication p-value and partitioned into deciles; we then compute power to replicate the variants in each bin using effect estimates with or without the Winner’s Curse. Left panel (A): including all papers (WC-corrected *χ*^2^ goodness of fit *p* < 10^−4^); right panel (B): including only papers conducting discovery and replication in the same continental ancestry per variant and reporting accurate per-locus *N* (WC-corrected *χ*^2^ goodness of fit *p* = 0.67). Improvement of fit exceeds what is expected due to loss of power from subsetting data (*p* < 0.01). X-axis: upper p-value boundary of bin; Y-axis: predicted fraction of replication within corresponding bin based on power estimated from discovery data. Tracks correspond to predicted power to replicate using raw discovery (red) or WC-corrected (teal) effect estimates. Error bars correspond to 95% confidence intervals around mean replication rates as estimated across multiple loci.

### Functional Enrichment in Replicated Variants

Finally, we evaluated enrichment of functional annotations in detected and replicated variants. We restrict this analysis to 56 papers which imputed their discovery samples using the HapMap2 CEU reference panel. Variants in the CEU reference provide a null distribution for functional annotation. Amongst all 998 loci for which replication was attempted in these papers, the observed 29 nonsynonymous variants constitute 5X enrichment compared to expectation from HapMap2 (expected 6 loci; *p* < 0.0001 based on 10000 simulated resamplings of random variants matched on count and minor allele frequency). This is due to significant enrichment of genic SNPs amongst all replication candidates (3.6X, *p* < 0.0001), as well as an additional enrichment of nonsynonymous variants among them (1.5X, *p* = 0.0003). Variants reaching per-paper Bonferroni replication are further 1.8X enriched in nonsynonymous exonic variants, from 2.9% across 998 attempted variants to 5.2% in 443 replicated ones (Binomial test one-tailed *p* = 0.0061). This change is due to enrichment of exonic SNPs in replicated variants, with no further significant selection for functional variants (*p* = 0.37). These results are not being driven by particular outliers (*χ*^2^ goodness of fit *p* = 0.44; Methods). Analogous enrichment among nominally-replicated variants (1.3X) is not significant (Binomial one-tailed *p* = 0.1447).

## Discussion

This study provides the first systematic evidence of the efficacy of internal replication in the field of quantitative trait genome-wide association studies. Overall, with important caveats, we find that the field as a whole publishes results that replicate in a manner consistent with their expected power to replicate; this seemingly argues against the possibility of systematic flaws in GWAS methodology. The two significant predictors of aberrant replication performance, beyond the Winner’s Curse itself, are (1) incorrectly reporting maximum sample size instead of per-variant sample sizes, reflecting locus-specific missingness; and (2) conducting replication in samples of different continental ancestry than those used in discovery. Corresponding to reporting error and linkage disequilibrium effects, these influences are not surprising. Yet we have shown (S6 Fig, S7 Fig) that, within the papers considered here, these factors are necessary and sufficient to explain all internal replication discrepancies. This result is both novel and reassuring.

Though we present data separately for papers violating one of the two consistency conditions (see S9 Fig, S10 Fig, S3 Table), we do not present extensive analysis or conclusions for these substrata. Unfortunately, the number of papers in each bin becomes quite small, in particular the mere four papers with different ancestries in discovery and replication but correct per-locus sample sizes. With such small counts, given the large paper-level heterogeneity we observe in Fig 1, we hesitate to draw conclusions about these subsets. This prevents direct evaluation of the relative importance of ancestry and sample size in replication prediction; in practice, it is likely variable, dependent on the rate of missingness in a given study and the relative divergence between the ancestries considered.

Several strategies have been developed for accounting for the Winner’s Curse in reporting of signals. The use of multiple stage GWAS, in which samples are conceptually partitioned into (possibly several) “discovery” and “replication” phases for internal replication, may be considered an attempt at removing positive bias in effect estimates. The discovery samples are used to reduce the pool of candidate SNPs from ~10^6^ to ~10^1^ –10^3^, at which point replication samples are used to verify that the selected SNPs maintain their direction and approximate magnitude of effect in an independent sample. Unfortunately, in many studies that make use of the discovery and replication partition, the final reported results are not solely based on the replication sample. Most commonly citing the argument in [17], studies frequently meta-analyze effect estimates from discovery and replication for a given SNP. This joint estimate maintains the benefits of prioritization by discovery, namely in reducing the cost of the study by minimizing the number of variants assayed in the replication samples. However, this estimate incorporates the probabilistically biased estimate from discovery, possibly attenuated by the less-biased estimate from replication. Thus while the argument of [17] holds, stating that meta-analysis of two-stage studies maximizes power to detect variants, this increase in power comes at the cost of both increased false positive rate and significant bias in the estimate of effect at true, detected signals.

Our selection of the Winner’s Curse correction method of Zhong and Prentice [3] is based on two considerations. First, and most importantly, we lack access to the raw genotype data behind the loci we consider, as is required in [4]. Moreover, the number of variants reported in each study is unpredictable: in some cases there is just a single variant reported, whereas in others the investigators considered several hundred. In the case of the former, we cannot reliably apply methods like [18], which intrinsically require summary data from a large number of the most associated variants in a study to generate an effect estimate distribution. Both of these limitations strongly call for a method such as [3], in which correction is applied individually to each variant using only summary data. Yet this remains a limitation of our study, as we are not practically able to evaluate alternate methods of WC correction.

This study exclusively addresses the performance of internal replication in quantitative trait GWAS. Yet other forms of replication are of just as much importance to the field. External replication, in which the results of one study are tested by independent investigators, is an important metric of the reliability of a field. In other contexts, external replication is known to perform at much lower rates than internal replication, suggesting various forms of bias. There is furthermore the consideration of functional replication, here broadly meaning the extent to which meaningful biological insights can be derived from GWAS data. Both of these forms of replication are largely unaddressed by the current study, nor indeed are they considered by the majority of GWAS publications; this does not diminish their importance. It is entirely possible that our results concerning the correct performance of internal replication may coexist with extremely low rates of external replication. Yet internal replication itself remains an important component of the field, and one in need of proper characterization. This study emerged from a discussion of the performance of internal replication in GWAS, when we discovered that there were no available data to prove one way or the other whether the internal replication model was effective in practice. We hope that our analysis provides one measure of reassurance about the fundamental reliability of the GWAS model.

Perhaps the most unusual observation of this analysis is the substantial proportion of manuscripts in the field that do not provide enough information to actually allow independent validation of their results. While some of the filters applied in our QC pipeline were present simply for ease of modeling, at least 58% of papers we collected failed to include the minimal amount of reporting to fully prove their claims of replication. This situation is a failure both in data reporting by authors and by peer review in journals. Combined with variable definitions of replication, we suggest this accidental lack of transparency substantially contributes to perceived unreliability of statistical genetics within other scientific disciplines. A higher standard of reporting, that will not only enable computation of unbiased effect estimates, but also list them explicitly, may be beneficial for the field.

We detect significant evidence for publication bias, the preferential publication of results based on their perceived quality. In particular, as shown in Fig 1, journals of higher impact factor, most notably Nature Genetics, published studies that replicated more variants than expected based on their statistical power; the inverse relationship is true as well. While this may intuitively suggest that the most robust results are published in the best journals, there is no intrinsic reason that the results of a study that replicates according to its statistical power distribution should be less robust than those of a study that replicates more often than should be possible. Rather, as is often the case with publication bias in other contexts, we raise concerns that the competitive publication of GWAS is giving rise to a publication record with invalid statistical properties, an important consideration that is not widely appreciated at this time.

Somewhat surprising is the lack of a clear ranking bias, in which studies combine variants at a locus according to strength of association, thus biasing each locus’s indicator SNP beyond the standard Winner’s Curse. This process, which has been termed “LD clumping,” is reasonably common, but was not consistently reported as used in the papers we analyzed. In some cases, papers reported data for all variants at a locus, and we implemented our own version of LD clumping, by randomly selecting a variant at each locus and discarding all other variants within a certain conservative physical distance. This process may have somewhat attenuated any ranking bias in this dataset; but it is quite likely that some of the fit deviation observed in our dataset is attributable to ranking bias, yet is simply not strong enough to create systematic significant deviation at the granularity of the tests we apply here.

The indirect method of data collection used in this study raises several difficult questions concerning data consistency. Due to the sheer volume of papers analyzed in the course of this study, we must assume some errors are included within our data: both in the form of flawed data collection by the authors of this paper, and mistaken reporting from the individual papers that was missed in both peer review and our manual inspection. Of particular note, for several tests included in this study, we have assumed for our statistical models that these papers report complete sets of loci brought to replication. We have furthermore specifically removed papers that transparently report partial subsets of results. However, without access to raw SNP lists from the contributing GWAS, there is no method to directly verify this criterion. There are also concerns about the low precision of data typically reported in GWAS publications. While we are able to make bulk conclusions across many loci, calculations at individual loci are somewhat unreliable. In particular, we have attempted to recompute the apparent sample size per variant for studies that have reported maximum sample size only; however, low precision data have made the resulting sample sizes rather unreliable, even generating in many cases sample sizes larger than the original value. Of note, since we cannot rescue the replication predictions in papers with different replication ancestry or maximum sample size reporting, there remains the possibility that sample size reporting or different ancestry is merely a proxy for a more meaningful underlying set of variables that selectively impacts papers with these apparent reporting flaws. Future analyses of this kind would strongly benefit from access to more of the raw data from contributing studies, should the resources be available for such an undertaking.

It is important to note that the restriction of analysis to quantitative (normally distributed) traits limits the direct conclusions we may draw to those same studies. This leaves the remainder of GWAS, which typically study binary disease traits and make up approximately 67% of the NHGRI-EBI GWAS Catalog. The methods used in [3] were initially developed for case/control studies and operate on regression test statistics, meaning the approach we have taken here may be easily applied to case/control studies in a later analysis. However, the practicalities of data collection meant that it was not possible to more than double our data acquisition for this study. We see no particular reason to assume different conclusions will be drawn based on binary trait studies, and suggest that our conclusions may provide a reasonable starting point for the interested analyst.

This study is not designed to counter-productively single out individual papers or investigators. For transparency, the full citation list is included (S1 File). We directly disclose our own statistics among considered papers. I.P. did not author any; C.P. contributed to 12 papers (3.6%) in the initial pool, two passing QC (consistent with expectation, Binomial given overall rejection rate across all papers, *p* = 0.7751). Seven of the ten removed papers provided incomplete data for replication, more than expected by chance (Binomial given rate of this error across all papers, *p* = 0.007). This anecdotal observation of papers focusing on anthropometric traits suggests the consistency of stylistic conventions within a phenotypic field to translate into recurrent faults in data reporting.

## Conclusion

The Winner’s Curse correction algorithm used here is based on a simple and fast method of generating unbiased effect estimates [3]. Our implementation [19] requires simple input parameters (replication threshold, SNP frequency, etc.) available from studies in the field with no paper-specific modifications required. This tool models a traditional two-stage GWAS design, as opposed to a paradigm of merging data from both study stages [17]. While strict staging is less powerful in detecting true associations, meta-analyzing discovery and replication results in effect estimates still subject to directional bias from discovery, and is thus not considered in our software.

This analysis provides the first systematic evidence that quantitative trait association studies as a whole are replicable at expected rates. The fairly lenient quality control required to generate such a result is instructive: papers conducting discovery and replication in populations of similar ancestry and reporting accurate sample sizes replicate according to their predicted power. That these criteria are met in only 12% of all successfully published papers indicates intrinsic flaws, not in the paradigm of GWAS, but rather in study design and reporting standards. Correction of discovery effects provides distinct advantages for any GWAS study. Most fundamentally, replication at expected level is a sanity check for the analyst. Furthermore, WC-correction allows rational and optimal prioritization of variants for replication. Finally, as a field, it is critical for GWAS to report correct, rather than inflated results.

## Materials and Methods

### Notation

We consider *M* independent loci brought forward to replication from all papers combined. Each individual paper x contributes *M_x_* loci to this total. A variant has an estimate of effect *β_obs_* on a given phenotype as well as a standard error of that estimate *s*, both computed from some form of linear regression. Significance, either for bringing variants forward from discovery, or for considering variants successfully replicated, is defined based on a p-value threshold *α_x_.* The corresponding test statistic 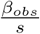 is standard normally distributed; *ϕ, ψ* are thus the PDF and CDF of the standard normal distribution, respectively.

### Data Collection

The NHGRI-EBI GWAS Catalog [1, 10] is an online resource that collects certain annotations for all SNPs reported as significantly associated with a human trait. As significant association and successful peer review are the only major criteria used for inclusion in this database, we used it as a reasonably unbiased source of papers in the field across a variety of phenotypes and journals. We restricted the articles selected from the database to fit our modeling requirements as follows. The papers selected must primarily:

1. study at least one quantitative trait;
2. be published in a journal with a primary focus on human genetics;
3. provide regression effect, standard error, frequency, and sample size for both discovery and replication;
4. provide data for all variants brought forward to replication; and
5. model a minimally two-stage (discovery and replication) study design with a p-value threshold used to select variants for replication.

The full list of filters and papers lost due to each criterion is shown in S2 Table. Whenever possible, we made reasonable accommodations to the papers to attempt to include them in this study. We consider variants novelly discovered in each paper, as opposed to those previously reported for a trait in question, as those are the variants typically brought forward for replication. Papers conducting multiple GWAS (i.e., reporting multiple phenotypes tested in the same study sample) had all novel discovered variants from all traits included in the analysis, and are conservatively reported as a single unit in this analysis. For studies that reported a single allele frequency per variant, as opposed to a distinct frequency for each of discovery and replication stages, we used that one frequency for both stage instead. Studies that did not report a variant-specific sample size, to accommodate for differential missingness at different sites, were assigned the maximum available sample size assuming no per-site missingness. These modifications will introduce noise into the final analysis, yet a large percentage of papers required at least one of these modifications and thus were permitted in the interest of representation and sufficient sample size.

Studies with different replication designs were compelled whenever possible into the traditional two-stage format we use here. Thus for studies that attempted multiple non-tiered replications, followed by a meta-analysis of all discovery and replication panels together, we conducted the replication study meta-analysis manually using standard error weighting in METAL [20]. Studies that conducted tiered replications were included with the first tier replication, in which all variants passing a threshold from discovery were tested, used for their replication study.

### Winner’s Curse Correction

To perform bias estimation, we use an implementation of the model in another study [3]. The major benefit of this model is that it may be applied to variant summary statistics as opposed to raw genetic data. As the non-parametric method BRsquared [4] requires raw genetic data, we did not consider this alternative. The maximum likelihood model we use is as follows:

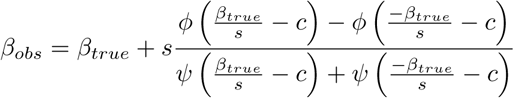

Here, *β_obs_* is the (likely biased) effect estimate observed in discovery; *β_true_* is the conceptual underlying unbiased effect of the variant in the source population; and *c* is the test statistic corresponding to the discovery *α* threshold in a given study. The expected bias of the observed effect, *E*[*β_obs_ − β_true_*], scales inversely with the distance between the observed test statistic and the cutoff applied to variants brought forward to replication. The bias can be solved using any standard zero-finding algorithm (for example, Brent’s method as implemented in C [21]). Note that in situations in which the observed test statistic far exceeds the *α* threshold, each component of the bias in the above equation is dominated by one or the other of the paired terms; only when the statistic is close to the threshold (that is, when the expected bias is large) do both terms meaningfully contribute to the bias estimate.

### Independence of Loci

To simplify predictions of replication efficiency, we considered an independent subset of all reported loci. As we lack direct access to the genetic samples used in these studies, we extracted a subset of the variants such that no two variants in a paper are situated within one megabase of any other. This is a very simple modification of the standard clumping protocol used in GWAS studies [22]. To prevent additional bias, we report a random variant from each locus, not necessarily the most strongly associated in discovery. This will effectively guarantee that each variant represents a single locus with only minimal linkage disequilibrium between variants, but is conservative in the sense that it discards any secondary signals present among the replicated variants.

Furthermore, this approach may attenuate functional annotation burden testing if the strongest association in an LD block is preferentially causal. While certain papers specifically address the possibility of secondary signals by sequential conditional analysis of variants, the inconsistency of this analysis and absence of it in many papers led us to seek a uniform treatment of all papers in this study.

### Definition of Internal Replication

The concept of internal “replication” may be interpreted differently in different reports. We consider three definitions of replication for this study, to observe different characteristics of the data:

1. replication at nominal *α* = 0.05 [“nominal”]
2. replication at 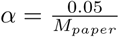 [“Bonferroni”]
3. replication within deciles of variants [ “deciles”]

We specifically only consider methods in which replication is determined from the replication study alone. The nominal and Bonferroni methods are commonly used. We use the decile method to investigate the behavior of the predicted power [23] to replicate according to the strength of an association signal. We compute decile goodness of fit with the Poisson binomial distribution, using average power to replicate in each bin across all variants. This permits a formal analysis of differential performance of replication at different levels of replication stringency.

Given a set of variants and their predicted power to replicate at a given *α* threshold, the number of observed replications is distributed as Poisson binomial with success probabilities equal to each individual variants’ power to replicate (see below). This is a generalization of a Binomial distribution in which each Bernoulli trial is allowed to have a known but variable success rate. We use the implementation of this distribution in R [24]. We further adapt the standard two-tailed Binomial test for use with the Poisson binomial CDF implemented in this package.

We note that under certain assumptions the number of replications will asymptotically be distributed normally. However, depending on the *α* considered, many variants analyzed here have power of effectively, or within machine precision, 0 or 1; with our limited sample size, the convergence properties of our dataset will be undesirable, and thus we use the exact distribution at the cost of computational efficiency. This process may be considered a fitting of the model according to which the WC-corrected discovery data correctly explain the observed replication data.

In several instances, we evaluate the effects of filtering certain subsets of papers based on various criteria, and the extent to which this causes fit criteria to return to null expectation. As this evaluation is potentially confounded by reduced statistical power, in all cases we test whether the change in p-value is significantly different from expectation under random subsampling of variants matched on total power to replicate amongst the observed variants.

### Power to Replicate

Assuming the discovery and replication sample of a study are drawn from the same source population with shared expected effect at each variant, the power to replicate a discovered variant *v* for a quantitative trait under the additive model is

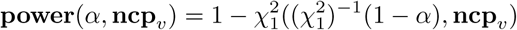

In brief, the power to detect a signal at an *α* threshold of *p* is the probability of the variant exceeding the required test statistic from the null, but under the alternative distribution which is noncentral 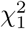 with per-variant noncentrality parameter

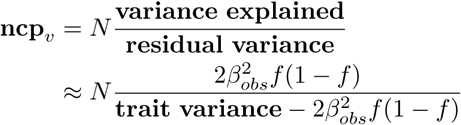

where *N* is the replication sample size and *f* is the replication allele frequency of the variant. This *f* should be the actual allele frequency in the replication sample; however, studies sometimes report *f_hapmap_* from the closest reference ancestry as a means of protecting patient anonymity. Overall, the predicted number of replications across all variants in a paper is the sum of the power to replicate, as a function of predicted effect size and replication sample size and frequency, across all variants analyzed.

### Sample Ancestries

These papers demonstrate the coverage of population ancestries in the field of quantitative trait genetics. We report and analyze the ancestral coverage of these studies using the simplifying summary statistic of continent of ancestry (Europe, Africa, East Asia), tracing generally the ancestries of the original HapMap2 populations. We include a fourth category for African American samples, the largest admixed population nonnegligibly represented in the papers. This geographical partitioning matches the ancestry assumptions used in GWAS methods such as genotype imputation.

Ancestry group counts are computed from maximum reported sample size per cohort per paper. In studies where cohorts of different continental ancestry are meta-analyzed, sample sizes are appropriately partitioned to the contributing ancestries. No adjustment is applied for papers reporting on the same cohort. For comparison to what the field’s sample sizes would be under random global sampling, global population estimates are computed [12,13].

### Functional Annotation

We tested loci for nonrandom annotations. This test is usually conducted with access to the full set of variants tested in an individual study. As in this study design such information is masked, we restricted the analysis to papers using HapMap2 imputation in their discovery data; considered only SNPs present in HapMap2; and restricted the data further to European ancestry discovery data, which includes the majority of papers in the dataset.

We annotated all variants in the CEU subset of HapMap2 using ANNOVAR [25]. We computed the average rate of functional annotations in the true set of variants. To generate a null distribution, we matched true variants on allele frequency and, when appropriate, whether the variant was located in an exon. P-values are computed over 10000 simulated null sets.

## Supporting Information

**S1 File Text supplement.** Contains: S1-S11 Figure, S1-S3 Table, and bibliography for all papers considered from the NHGRI-EBI GWAS Catalog.

**S1 Dataset Compressed archive of independent subset of loci, from all papers used in analysis.** Each file corresponds to a single PMID as listed in the filename. Only includes data for 100 studies used in analysis. For full citations, see S1 File.

**S2 Dataset Jupyter notebook of statistical analyses in the paper.** Included: the raw data from S1 Dataset, with per-variant data for each paper, split by ancestry and sample size consistency, both before and after Winner’s Curse correction; per-paper replication computations, both with nominal and Bonferroni replication; and Poisson binomial testing on per-variant decile data. Any other analyses from the study are straightforward to reconstruct from the variables and scripts in the notebook. To repeat the Winner’s Curse correction itself, use the C++ library release [19].

